# Whole-Genome Sequencing and Variant Discovery of *Citrus reticulata* ‘Kinnow’ from Pakistan

**DOI:** 10.1101/2022.12.07.519411

**Authors:** Sadia Jabeen, Rashid Saif, Rukhama Haq, Akbar Hayat, Shagufta Naz

**Affiliations:** Lahore College for Women University, Lahore, Pakistan; Decode Genomics, 323-D, Punjab University Employees Housing Scheme, Lahore, Pakistan; Citrus Research Institute, Sargodha, Pakistan

**Keywords:** *Citrus reticulata*, GATK4 pipeline, WG variant calling, Resequencing-‘Kinnow’

## Abstract

Citrus is a source of many nutritional and medicinal advantages, which is cultivated worldwide with major citrus groups of sweet oranges, mandarins, grapefruits, kumquats, lemons and limes. Pakistan produces all of its major citrus groups with mandarin (*Citrus reticulata*) being the prominent group that includes local commercial cultivars such as Feutral’s Early, Dancy, Honey and Kinnow. The present study was designed to understand the genetic architecture of this unique variety of *Citrus reticulata -*’Kinnow’. The whole-genome resequencing and variant calling was performed to map the genomic variability that might be responsible for its particular characteristics like taste, seededness, juice content, thickness of peel and its shelf-life. A total of 139,436,350 raw sequence reads using Illumina platform were generated with 20.9 Gb data in Fastq format having 98% effectiveness and 0.2% base call error rate. Overall, a total of 3,503,033 SNPs, 176,949 MNPs, 323,287 INS and 333,083 DEL were identified using GATK4 variant calling pipeline against *Citrus clementina* as a reference genome. Further, g:Profiler bioinformatics tool was applied for annotating the newly found variants, harbor genes/transcripts and their involved pathways. A total of 73,864 transcripts harbors 4,336,352 variants, most of the observed variants were predicted in non-coding regions and 1,009 transcripts were found well annotated by different databases. Out of total aforementioned transcripts, 588 involved in biological processes, 234 in molecular functions and 167 transcripts involved in cellular components in *Citrus reticulata*. In a nutshell, 18,153 high-impact variants and 216 genic-variants found in the current study which may be used for marker assisted breeding programs of ‘Kinnow’ to identify this particular variety among others and to propagate its valued traits to improve the contemporary citrus varieties as well.

**Graphical Abstract:** 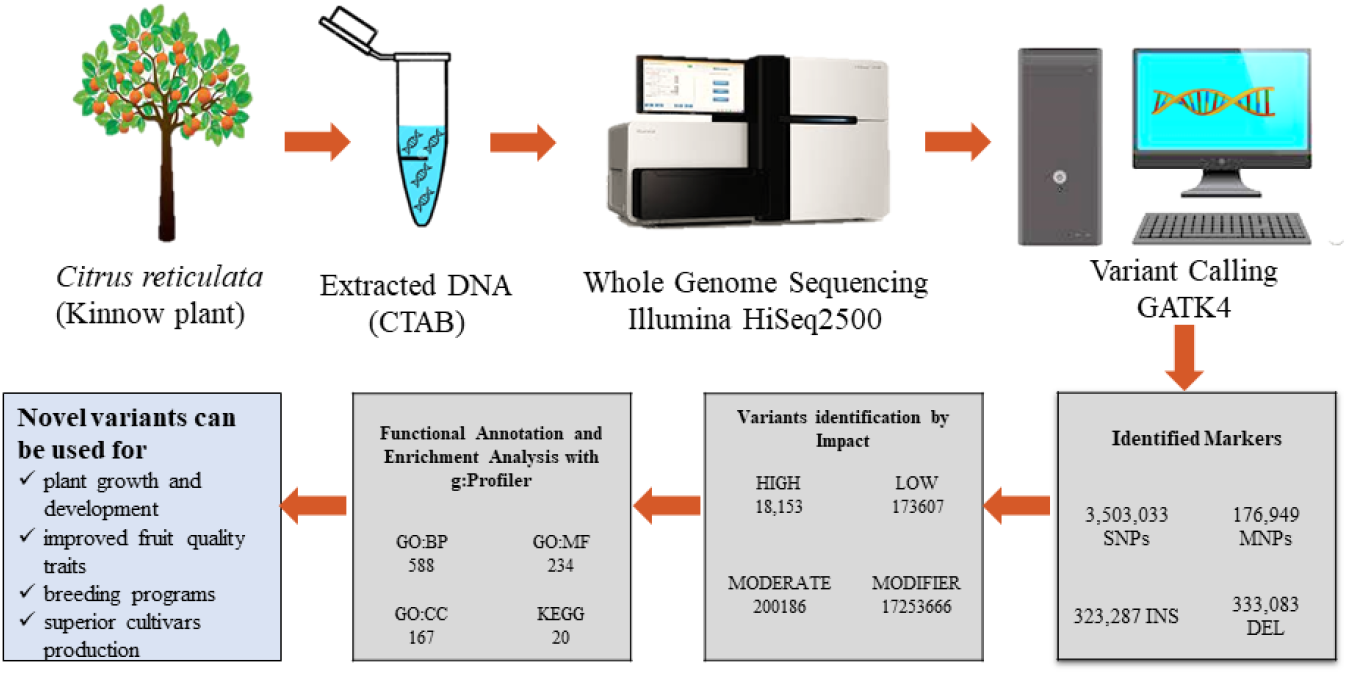

## Introduction

*Rutaceae* family comprising of 140 distinct genera and 1300 species including citrus genus that is cultivated globally (Kamal et al. 2011) and is rich in vitamins, minerals, carbohydrates, and dietary fibers (Prasad et al. 2015) holding all essential physicochemical properties (Hayat et al. 2017; Wu et al. 2018). It also possess other vital compounds such as “flavonoids” which are subject of anti-tumor and antioxidant properties (Etxeberria et al. 2009). It was given importance and was regarded as an excellent immunity booster during COVID-19 times. In 2019, world total citrus fruit production was of 158 million tonnes (Duru et al. 2022) while in 1970 it was just 39.6 million tonnes.

To date the annual production reported from Pakistan is of 2.29 million tons of Citrus fruits. Pakistan’s citrus fruit production increased from 445,000 tonnes in 1970 to 2.29 million tonnes in 2020, it is increasing at a 4.14 percent annual rate. This production covers an area of approx. 9,898,463 hectares with 115,554 hg per ha yield (Naqvi et al. 2022). *Citrus reticulata* ‘Kinnow’ is the lucrative fruit grown on 177.22 thousand hectares in Pakistan, with an annual production of 2116.47 thousand tones and it is important for country’s economy as accounting for about 97 percent of national revenue (Rehman et al. 2018). In the year 2019, Kinnow exports brought about $195 million in foreign exchange (Ahmed 2020)

There are different varieties of Kinnow cultivated in Pakistan including seeded and seedless (Rehman et al. 2018). Seedless fruit demand is high because of aesthetic reasons and ease in consumption (Zhao et al. 2020). The natural phenomenon behind seedless fruit production is parthenocarpy (Distefano et al. 2020). Pollen incompatibility, or ovule and pollen self-incompatibility are among the major natural causes of seedlessness in Kinnow mandarin (Altaf et al. 2014). Seedlessness is also linked with triploidy (Jaskani et al. 2005; Usman et al. 2008). In the absence of cross-pollination, male sterility avoids seed production and transfer of pollen grains to neighboring orchards. At the diploid level in citrus, several different types of male sterility related to chromosome aberration exist, resulting in varying degrees of pollen fertility (Khan 2007). Male sterility in citrus is a result of a gene–cytoplasmic interaction, as seen in satsuma, where male sterility is linked to pollen grain development failure and low viability (Goto et al. 2016). MS-P1, a large QTL for lowering the quantity of pollen grains per anther, and MS-F1, a QTL for decreased apparent pollen fertility, were recently discovered to be linked to male sterility (Montalt et al. 2021).

In this research *Citrus reticulata* (seedless) whole genome was compared with haploid *Citrus clementina* (seeded) as a reference genome to find genome wide genetic variations to have an insight of different characteristics of citrus fruits to promote marker assisted fruit quality and other valuable traits in future.

## Materials and Methods

### Identification and sample collection of true-to-type plant

True to type *Citrus reticulata* plant with seedless trait was identified with the help of senior research officers at Citrus Research Institute (CRI) (Fig. 1). This institute is located in Sargodha city of Pakistan which is famous for the production of quality fruits of citrus. This plant is purely a natural selection of seedless Kinnow from domestic orchard, selected by Niaz Ahmad, former Director CRI and planted in field in 2014 (Sabir 2010). For genomic DNA extraction leaves were collected and washed with distilled water followed by 70% Ethanol wash to avoid any kind of contamination and stored immediately in a cool box. In laboratory the mid rib was selected and chopped into small pieces to ease the grinding process in DNA extraction. Finally chopped leaves with proper labelling stored at -20 °C till further use.

**Fig. 1.**
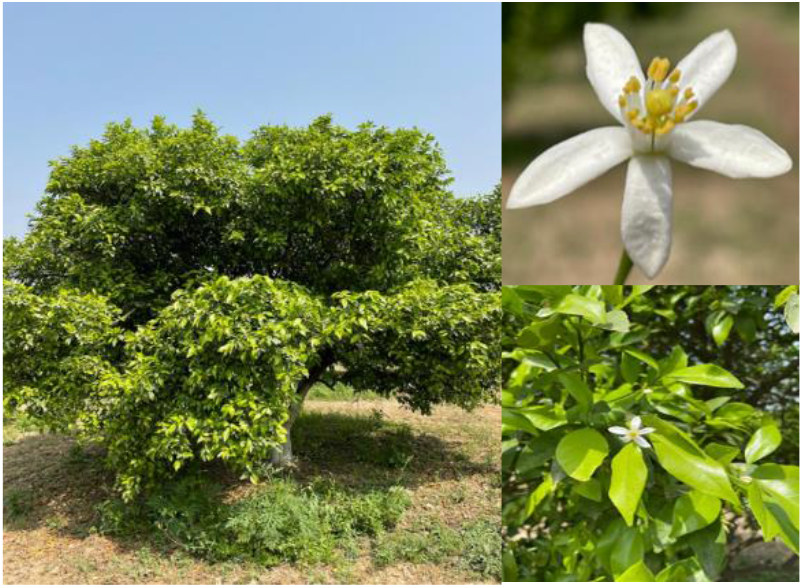
Seedless Kinnow plant used in the present study with branches and flower

### Genomic DNA extraction and quantification

The total genomic DNA of seedless citrus was extracted using the Murray and Thompson (Murray and Thompson 1980) protocol with some modifications (Naz et al. 2014). The frozen midrib material from leaves was ground into a fine powder in an ice-cold mortar and pestle. 1mL of extraction buffer CTAB (preheated) was added and incubated for half hour at 65°C. After this 500 μl of chloroform mixture was added to eppendorf tube, mixed it thoroughly and then centrifuged for 10 minutes at 6000 rpm. Supernatant was collected and transferred into new labeled eppendorf, again the chloroform mixture (24:1) was be added in an equal volume, and then centrifuged at 6000 rpm for 10 minutes. The same procedure was repeated by adding chilled propanol in an equal volume instead of chloroform. The addition of propanol caused precipitation of DNA molecule. The DNA pellet was resuspended in TE buffer (pH 8,1x) after being washed with 70% ethanol. The extracted DNA was stored at -20 °C. Nanodrop was used to check the purity and concentration of DNA. Before sequencing the DNA, samples were checked for contamination and DNA degradation by agarose gel electrophoresis (Sambrook et al. 1989, 2012). Further the DNA concentration was quantified by Qubit 2.0.

### Whole-Genome sequencing using Illumina platform

A total amount of 1.0μg DNA per sample was used as input material for the DNA sample preparations. Sequencing libraries were generated using NEBNext® DNA Library Prep Kit following manufacturer’s recommendations and indices were added to each sample. The genomic DNA is randomly fragmented to a size of 350bp by shearing, then DNA fragments were end polished, A-tailed, and ligated with the NEBNext adapter for Illumina sequencing, and further PCR enriched by P5 and indexed P7 oligos. The PCR products were purified (AMPure XP system) and resulting libraries were analyzed for size distribution by Agilent 2100 Bioanalyzer (Panaro et al. 2000) and quantified using real-time PCR. The qualified libraries are fed into Illumina HiSeq 2500 sequencer (Liu et al. 2022) after pooling according to its effective concentration and expected data volume of 20.9 GB paired end reads.

### Whole-genome variant calling bioinformatics pipeline

#### Quality Checks and Data Trimming

The raw reads after sequencing received in FASTQ format. At first the fastp tool was used for trimming and filtering the adaptors and low-quality reads from the FASTQ files. MultiQc was used to check the quality of reads after trimming (Hughes et al. 2022). It deleted the reads with a Phred quality score lower than Q30.

#### Mapping of filtered reads against reference genome

The fasta file and annotation file of reference genome *Citrus clementina (deploid)* was downloaded from (https://plants.ensembl.org/Citrus_clementina/Info/Index). Before mapping indexing of reference genome was performed by Bowtie2 and afterwards mapping of clean reads of our sample genome against index files of reference genome with the help of same tool Bowtie2 (Fernandez 2022). The resulting Sam file was converted into Bam file using samtools to conserve the space (Marbouty and Koszul 2022) *Variant Calling:* Before variant calling, we performed two steps AddOrReplaceReadGroups and Markduplications. Finally, the GATK was used for whole genome variant calling (Li and Durbin 2010). From the variant call format (VCF) files obtained through filtering, SNPs and InDels were annotated based on genomic location and classified by the likely effects of the variations, including functional categories using SnpEff (Cingolani et al. 2012; Naz et al. 2014).

#### Annotations

The SnpEff binary database file (.bin) was generated using the citrus genome annotation file (gff3) and the genome sequence. This generated database was used to annotate the effects of SNP by region effect (high, moderate, low, and modifier), and functional class (missense, nonsense, and silent) for all of the individuals’ scaffolds. The localization of SNP and InDel was based on the annotation of the gene models of the citrus reference genome (Wu et al. 2014). The two types of polymorphisms in the gene region and other genome regions were annotated as genic and intergenic, respectively. The genic SNP and InDel were classified as coding sequences (CDS), untranslated region (UTR), and intron, according to their localization. The SNP in the CDS were further separated into synonymous and non-synonymous amino substitution using SnpEff version (Jiang et al. 2022).

### Functional annotation and enrichment analysis

For the functional annotation and statistically significantly enriched terms of resulted SNPeff genes file, we used g:Profiler (https://biit.cs.ut.ee/gprofiler/gost). g:Profiler is a freely available web based tool provided four windows for gene profiling including g:GOSt, g:Convert, g:Ortho, and g:SNPense. g:GOSt analyzed individual or multiple gene lists for functional enrichment. g:Convert converts gene/protein identifiers between different namespaces. g:Orth allows to map orthologous genes across species. g:SNPense connects human SNP identifiers to genes. It has an advantage of presenting the results in the visual and statistical form. We uploaded the GO Ids of all the genes/transcripts with default parameters (Raudvere et al. 2019).

## Results

### Whole-genome sequencing data statistics

Whole genome sequencing was done from the extracted DNA of native variety of *Citrus reticulata*. The genomic DNA was randomly fragmented to a size of 350bp by shearing method, this fragmented DNA were fed into Illumina sequencer (Hi-seq 2500) and the resulted library DDSW200002244-1a were analyzed for size distribution on flow cell H7JYWCCX2_L1, H7YFGCCX2_L1, and H7JYLCCX2_L1. A total of 13,344,049 reads were generated on flow cell H7JYWCCX2_L1, 8,769,552 on H7YFGCCX2_L1, and 47,604,574 on H7JYLCCX2_L1. In total 20.9 Gb raw data in Fastq format were produced with 98% effectiveness and 0.2% base error rate.

### Quality checks of sequence reads

A total of 137,901,462 reads were passed the filters which are 98.9% of total reads represents the good quality of sequencing experiment. Out of total, 1,511,738 (1.1%) reads were removed due to low quality. Only 23,150 N sequences were obtained and excluded from the sequencing files. Also, 10.5% of duplication was observed in sequencing reads. Overall, high GC content as 38.5% observed.

### Genome-wide variant identification and annotation in *Citrus reticulata*

After indexing mapping of clean reads of *Citrus reticulata* against reference genome of *Citrus clementina* resulted in calling of 4,336,352 variants in total genome size of 301,364,702 bp, which indicates the occurrence of 1 variant after every 69 bases across the entire genome. Out of total aforementioned observed variants, 86925 multi-allelic VCF entries were also detected, similarly, 17,645,612 number of effects were noticed, which might be due to the fact that one effect may be consider as common for both sense and antisense strands. The average variant coverage was 4.31x in *Citrus reticulata*. This coverage is inconformity to the previous reported study in Papaya (Bohry et al. 2021). Around, 97.45% of newly reported variants harbor in first 9 larger scaffolds, while only 2.55% are present in remaining smaller scaffolds of approximately 17000, which seems comparatively conserved regions of the genome due to the fewer number of variants in these regions. Secondly, these markers may be more informative regarding the identification and other quality vested traits of this particular genome of *Citrus reticulata*. Current data is available under the NCBI SRA project ID: PRJNA821664.

Here, we highlighting only 23 variants which are observed in the CDS regions of the following genes e.g., FLA4, FLA12, FT1, NADP-ME3, ALDH7B4, CA2, XTH9, EXPA5, LAC1, AAO, 5MAT, AMY1, BAM1, CT-BMY, BMY5, BAM1, PHO1, PHS2, SEX1, PWD, DPE2, PTPKIS1, BMY3, AMY3 and GWD3. These mutations are supposed to have their effects in the genes functioning and ultimately their respective metabolic pathway in *Citrus reticulata*. These variants can be further screened with large number of samples to identify and propagate the valued traits to other varieties of the citrus by adopting targeted marker-assisted breeding strategies. The above-mentioned variants represented in Table 1, while remaining variants in the same genes is provided as Table S1.

**Table 1.** Whole-genome variants identified in the coding regions

| Gene/Function | Position | Variants | Type |
| --- | --- | --- | --- |
| FLA4/Cell wall proteins | 5311778 | C>T | SNP |
| FLA12/Cell wall proteins | 22185231 | T<G | SNP |
| FT1/Hemicellulose | 7416255 | C > T | SNP |
| NADP-ME3TCA | 7405816 | C>T | SNP |
| ALDH7B4/TCA | 7265446 | T>C | SNP |
| CA2/TCA | 22676285 | T>C | SNP |
| XTH9/Cell Wall Modifications | 21059179 | A>G | SNP |
| EXPA5/Cell Wall Modification | 25035219 | AT>A | Del |
| LAC11/Simple Phenols | 1876879 | G>T | SNP |
| AAO/Simple Phenols | 8436872 | A>G | SNP |
| 5MAT/Flavonoids | 27394538 | C>G | SNP |
| AMY1/Degradation Starch | 40191656 | G>T | SNP |
| BAM1/Degradation Starch | 7580175 | C>G | SNP |
| CT-BMY/Degradation Starch | 4633734 | G>T | SNP |
| BMY5/Degradation Starch | 21319244 | T>G | SNP |
| BAM1/Degradation Starch | 695273 | C>T | SNP |
| PHO1/Degradation Starch | 25331966 | C>T | SNP |
| SEX1/Degradation Starch | 22402305 | CA>GG | MNP |
| PWD/Degradation Starch | 18155181 | G>A | SNP |
| DPE2/Degradation Starch | 21768348 | G>A | SNP |
| PTPKIS1/Degradation Starch | 7413952 | T>TGC | INS |
| AMY3/Degradation Starch | 23578390 | C>T | SNP |
| GWD3/Degradation Starch | 28860169 | C>T | SNP |

Maximum number of variants were identified from the scaffold-3 with 792,719 (18%) of the total observed variants, followed by scaffold-5 with 723,410 (17%), scaffold-9 with 637820 (15%), scaffol-2 with 490,723 (11%), scaffold-8 with 399,204 (9%), scaffold-1 with 312,044 (7%), scaffold-6 with 311,595 (∼7%), scaffold-4 with 297,566 (∼7%), scaffold-7 with 260,785 (6%) and remaining scaffolds with 110,486 (∼3%) (Fig. 2).

**Fig. 2.**
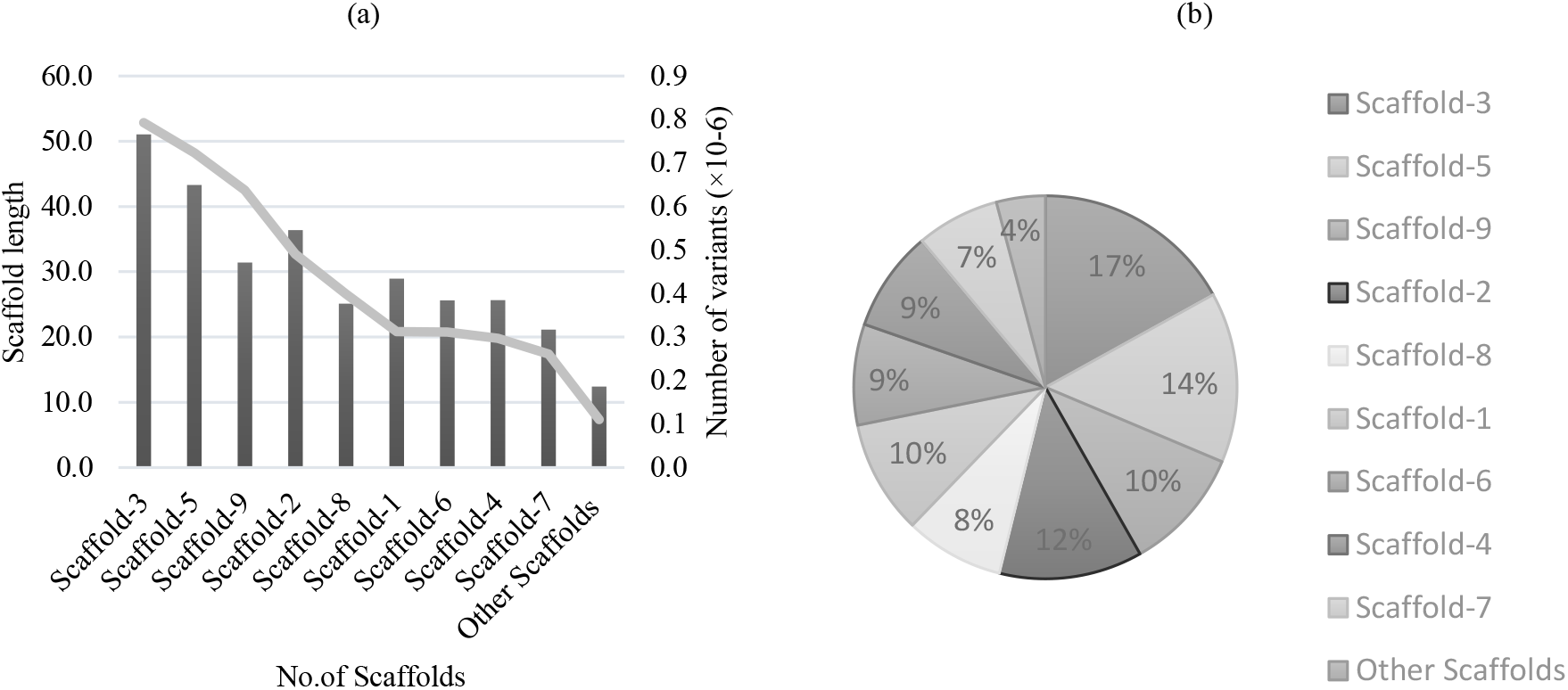
Localization and numerical total of observed variants in the genome scaffolds (a) Scaffold length (Mb, line) and number of observed variants (b) percentage of variations in each scaffold

### Categorization of variant by impact

Variants impacts sorted into four main categories: (i) high (ii) moderate (iii) low and (iv) modifier. High impact variants directly affect the gene and resulted in the loss of protein function. Variants caused the high impact with most common as stop-codon gained, stop-codon lost and frameshift variants, which may cause truncation of protein. Moderate impact variants are non-disruptive that normally change protein effectiveness, among moderate variants the most common found in our genome are missense and in-frame insertion and deletion (non-synonymous substitution) which causes amino acid changes in protein. Low impact variants are assumed to be harmless as it causes no change in amino acid residues, ultimately protein behavior (Synonymous variants). Modifier variants effects the non-coding region of gene like 5’ and 3’ UTRs, intergenic regions, and introns in genes (Fig. 3).

**Fig. 3.**
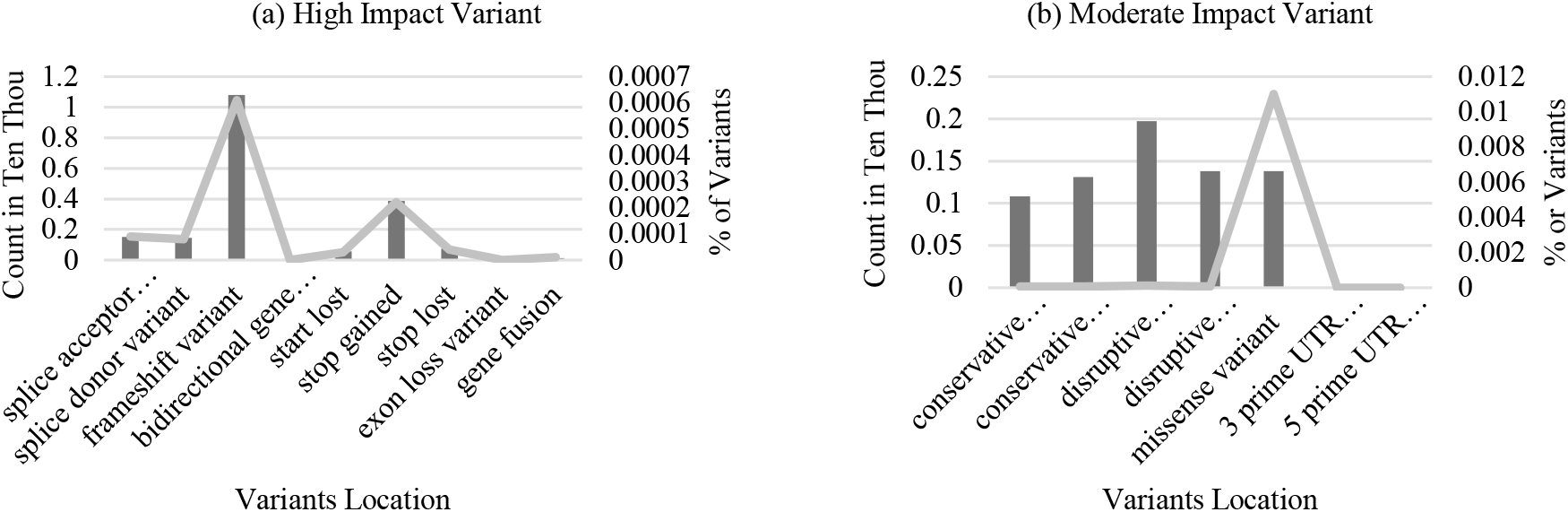
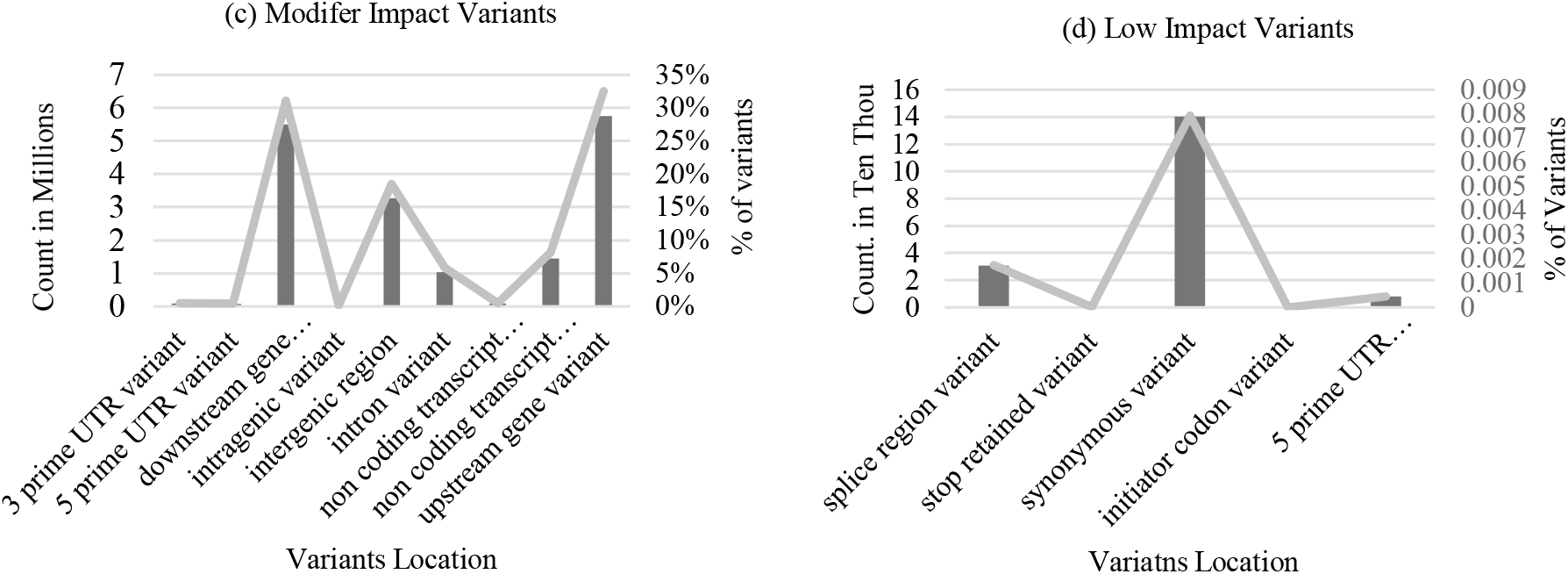
Illustration of variant categories by its impact on genome regions (a) High impact variant (b) Moderate impact variant (c) Modifier impact variant (d) low impact variant

### Distribution of whole-genome variants by genomic region

The reported variants were found in the different landscape of the genome including introns, exons, upstream, downstream, splice sites, UTRs, genic and intergenic regions. The most of the variants were observed in the regions of upstream and downstream with 5,751,269 (∼33%) and 5,493,167 (∼31%) counts respectively. While the other variants reported from the intergenic regions with 3,275,685 (∼19%), from transcript regions with 1472867 (∼8%), from intronic and exonic regions with 1,009,332 (∼6%) and 438657 (∼2%) counts respectively. Remaining regions like 3’ & 5’ UTR, splice site regions, splice site acceptor and donor harbor less variants as compared to aforementioned regions. Only 216 variants were observed from genic landscape which probably indicate the atmospheric adaptation of *Citrus reticulata* genome in Pakistani temperate geographical location. Graphical illustration of different variants are represented in the Fig. 4.

**Fig. 4.**
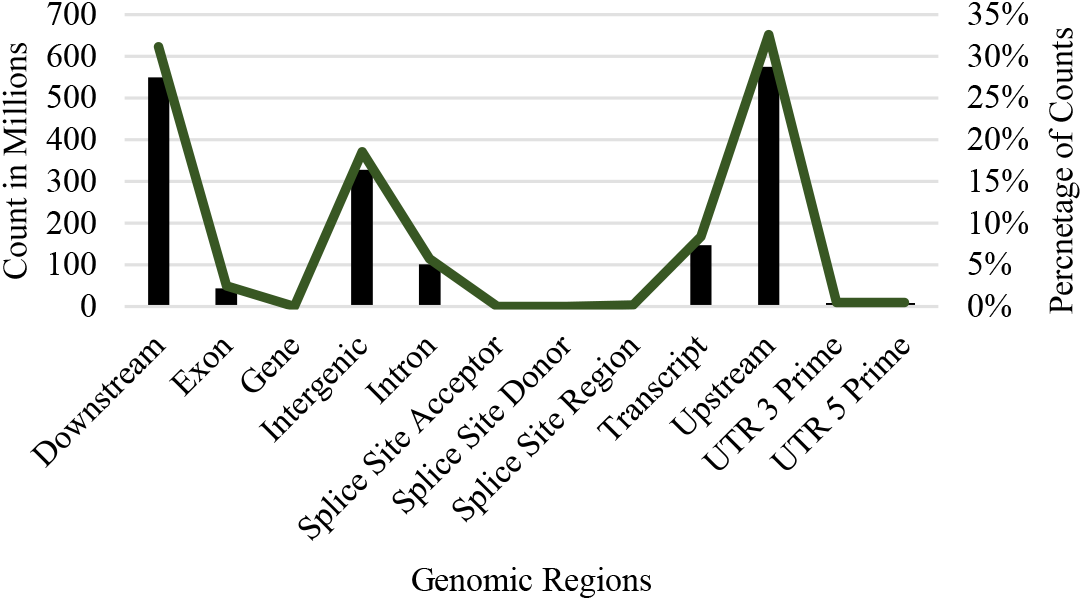
Graphical distribution of WG variants by genomic region *y axis numeric represents count in millions

### Distribution of WG variants by its type

The newly found variants from whole genome of *Citrus reticulata* were fall into four different categories/types of potential markers as Single Nucleotide Polymorphism (SNPs), Multi-Nucleotide Polymorphism (MNPs), Insertion and Deletion (InDels). Out of total 4,336,352 variants, the maximum numbers were identified as SNPs 3,503,033 i.e., the ∼80% of the total variants. The other class of identified variants was InDels, which is the ∼15% of the total variants, (INS are 323,287 (7.45%) and DEL are 333,083 (7.68%)). Remaining 176,949 are MNPs which are the ∼5% of the total variants (A variant is counted as variant when the reference allele 1 and allele 2 are all an equal length other than 1). Comparatively, SNPs marker is the most frequent in our studied genome due the simplest nature of point mutation, InDels markers may interfere in the physiological, biochemical and other pathways of the plants so observed in moderate count. On the other hand, MNPs are considered as critical markers and observed in lesser count due to its complex nature. Fig. 5 is the graphical representation of distribution of variants with its percentages.

**Fig. 5.**
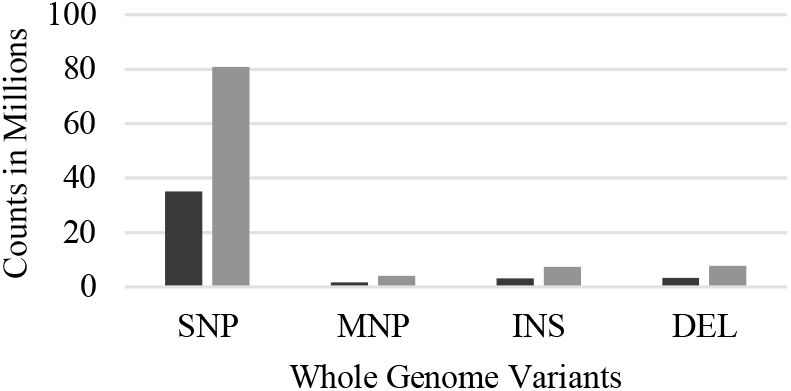
Distribution of variants on the basis of class. Black and grey bar shows observed counts and percentage respectively.

### Identification of number of variants by functional class

Critical information for building and maintaining the living organism is warehoused not only in the genetic code but also in the regulatory sequences adjacent to, within, and distal to genes (DNA). This is translated into functional proteins using the genetic code which consists of three nucleotides, and the correct order of these nucleotides is crucial as it encodes a specific amino acid sequence in protein. The changes in the arrangement of nucleotides resulted in the mutations which are categorized as missense, non-sense and silent mutations. In the current study genome 181,595 missense mutations were observed. While in the nonsense mutations the swapping of single nucleotide resulted in the production of stop codon, thereby translation stop prematurely as a result truncated protein produced, that may or may not be functionally affected. Only 3,183 nonsense mutations found in genome of *Citrus reticulata*. The last category i.e., the silent mutations, do not alter the sequence of amino acid but can still influence splicing accuracy or efficiency of the genome. 139,977 silent mutations were observed in the present study genome. Fig. 5 is the graphical representation of classes of the variant’s effects as missense, nonsense, and silent.

**Fig. 5.**
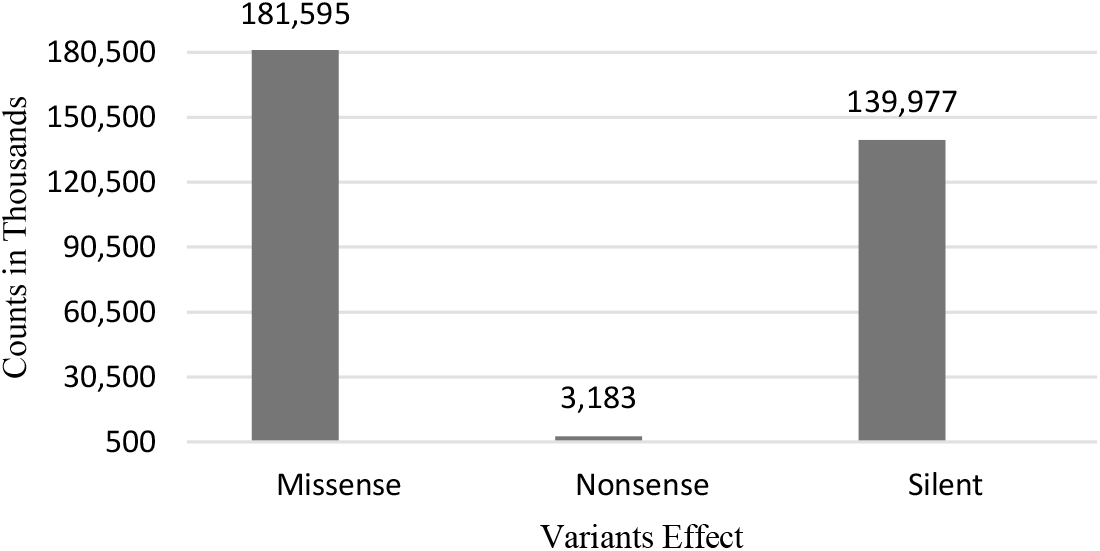
Classes of the variant’s effects

### Transitions & Transversion ratio in *Citrus reticulata*

Transition mutations normally higher as compared to the transversion mutations due to spontaneous tautomeric shifts. The current results of WG variant calling shows that the most commonly observed variants were transitions in purine and pyrimidine bases, Guanine (purine) replaced the Adenine (purine) by 616,615 times and the Cytosine (Pyrimidine) replaced the Thymine (Pyrimidine) by 614,000 times. Further observations indicate the following changes Adenine replaced the Guanine by 523719 times, Thymine replaced the Cytosine by 523301 times, Adenine replaced the Thymine by 188,683 times, Thymine replaced the Adenine by 188,353 times, Guanine replaced the Thymine 171,120 times, Cytosine replaced the Adenine by 170,534 times, Adenine replaced the 155,568 times, Thymine replaced the Guanine by 155,295 times, Cytosine replaced the Guanine by 98274 times, and the lowest changes observed as transition i.e. Guanine (purine) replaced the Cytosine (pyrimidine) by 97,571 times. Fig. 6 is the graphical representation of base change observed in the present genome. Collectively we found 2,242,354 transitions (Ts) and 1,183,807 transversions (Tv) mutations, with a Ts/Tv ratio of 1.8 (Fig. 6b and 6c).

**Fig. 6.**
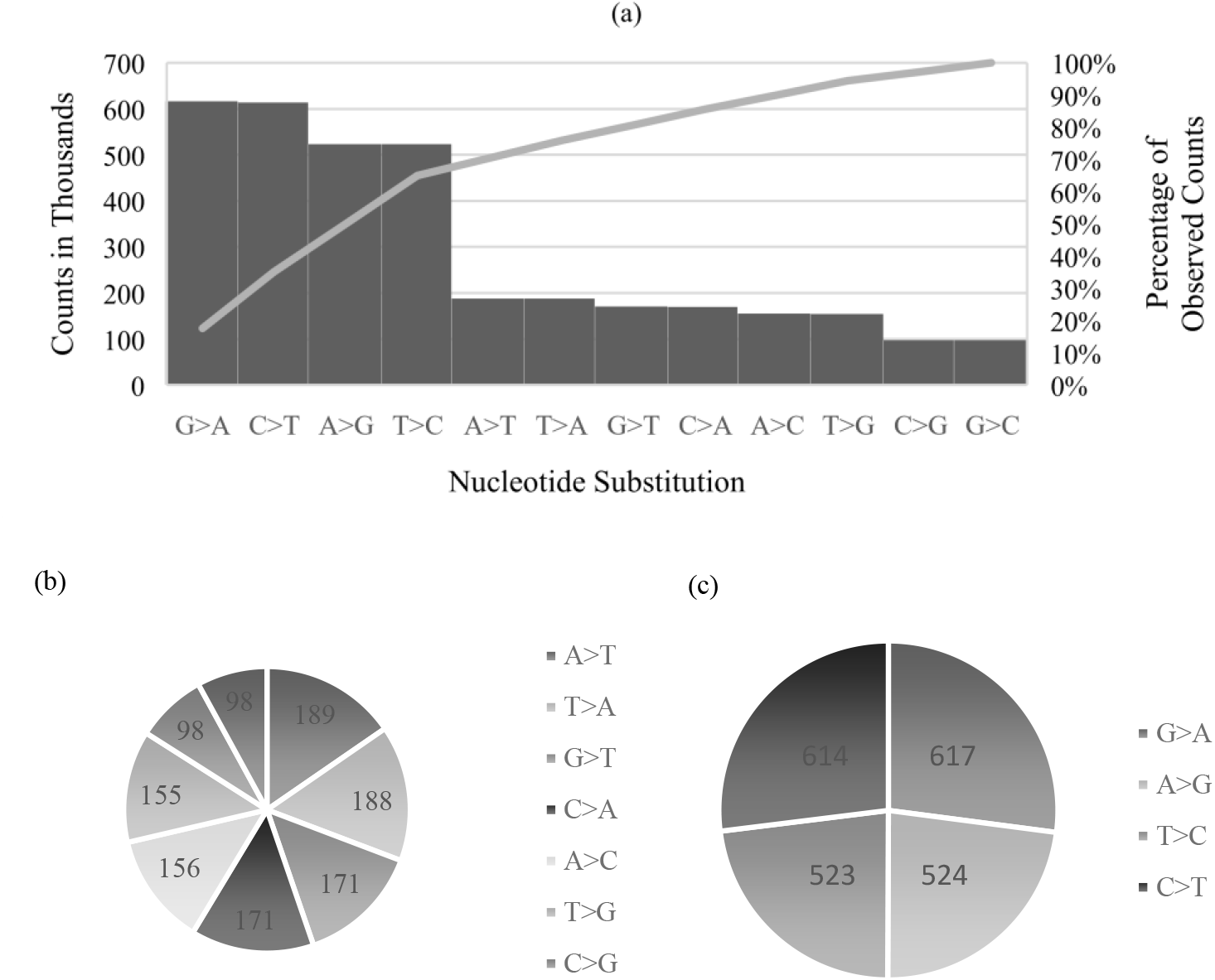
(a) Transition and transversion frequency with percentage nucleotide substitution in genome (b) Transversion changes (c) transition changes

### Predicted genome-wide codon changes

The genetic changes at DNA level ultimately alter the protein structure and eventually its function in different biological processes that may or may not change the physiology of living organisms. The alterations in whole genome of *Citrus reticulata* also predicted some changes at codon level due to the SNPs, InDels, MNPs and frameshifts. In total, 112, 906 codon changes were examined in analysis. Highest codon change was observed as GAC>GAT (3010 times), GAT>GAC (2962 times) and TCG>TCA (2703 times), AAC>AAT (2601 times), AAG>AAA (2591 times), GAG>GAA (2524times), GAG>GAA (2524 times), TCA>TCG (2515 times), AAT>AAC (2440 times) and CCG>CCA (2433 times). Details of complete codon changes are given in Table S2.

All the essential amino acids were predicted in the present genome which are accountable for plants cell division, pigmentation, production of natural hormones such ethylene, auxin and gibberellin. Fig. 7 represents the whole codon change in citrus genome

**Fig. 7.**
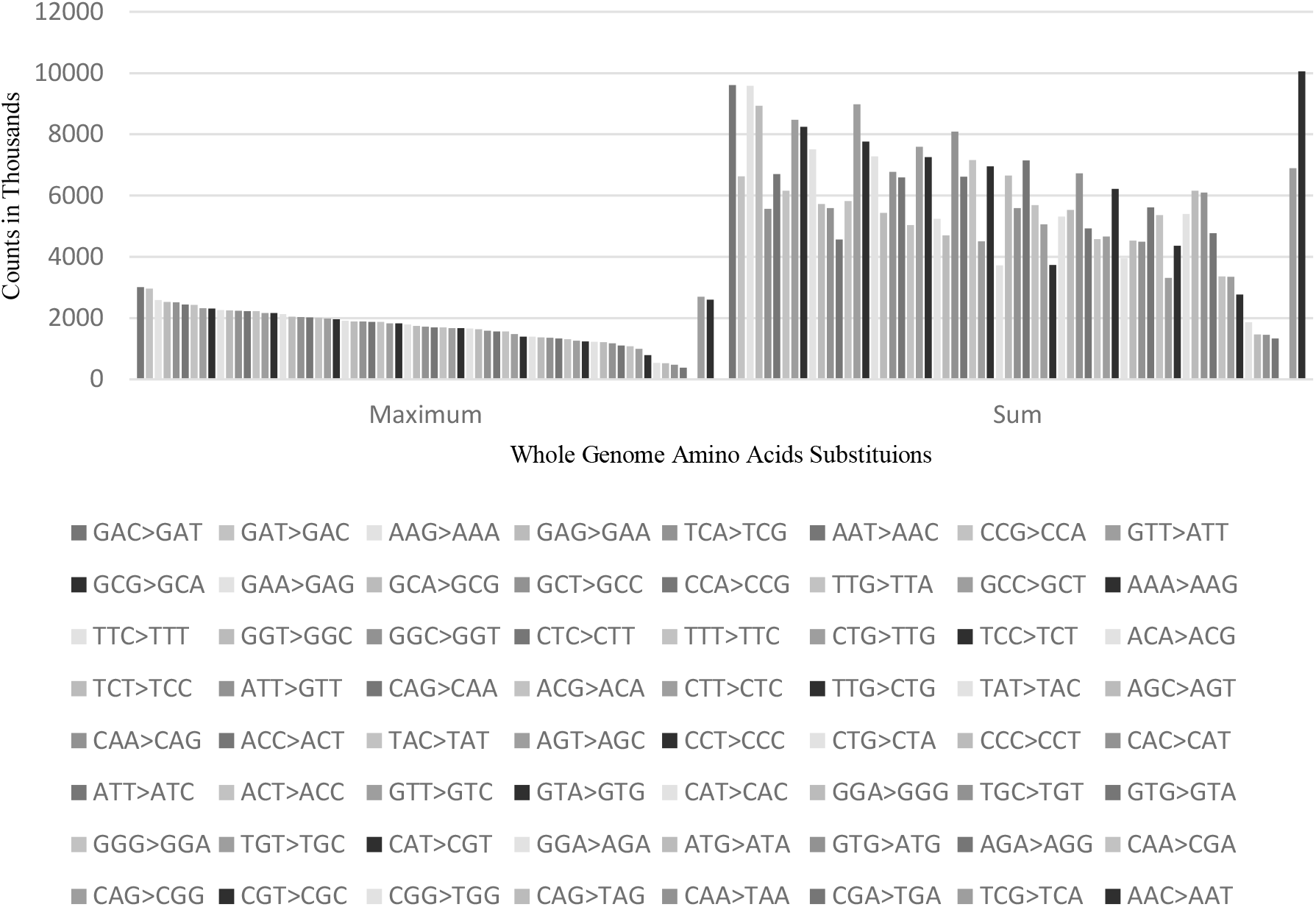
Predicted codon changes result of variations

### Functional correlation of newly found variants

In total 73,862 variants in our studied genome of *Citrus reticulata* were annotated from g: Profiler platform, this online tool compared our reported variants with different databases, as a results 1009 predicted transcripts were qualified on the basis of well described annotations. Based on adjusted *p-value*, 588 transcripts belongs to different genes involved in biological processes (GO:BP), 234 transcripts identified to have their roles in molecular functions (GO:MF), while 167 transcripts found to be associated to form the cellular components (GO:CC) and 20 transcripts belongs to metabolic pathways of the *Citrus reticulata* (Fig. 8).

**Fig. 8.**
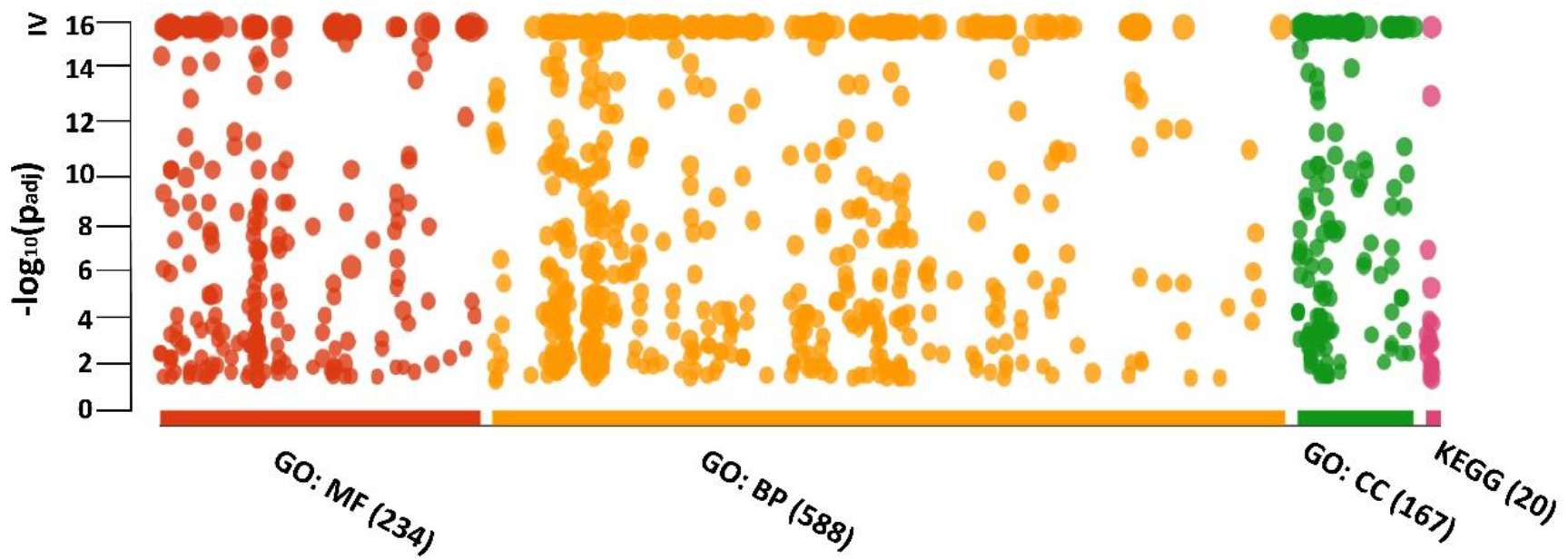
Graphical illustration of functional annotation of whole genome predicted transcripts

Our observed genomic variations were mostly placed in the transcripts predicted to function in to biological processes which indicates the physiological and homeostasis variability of this fruit plant. Organic cyclic compound metabolic processes in GO:BP were observed as top most on the basis of annotations from different databases (Fig. S1). Oxidoreductase activities in GO:MF (Fig. S2), while intracellular anatomical structures in GO:CC (Fig. S3). Ribosome, amino acid biosynthesis and glutathione metabolism were identified as top most among pathways (Fig. S4). Further functional annotation of whole genome transcripts is given in Table S3.

## Discussion

Pakistan is an agriculture-based economy comprises many of its subsidiaries including animal, crops, horticulture, pomology, forestry and fruit sectors. Fresh fruit industry played a pivotal role in revenue generation, particularly the *Citrus reticulata* (Seedless Kinnow) which is 30% of country total fruit output. Pakistan generated its highest revenue of $∼300,000,000 by exporting 460,000, 000 kgs in 2021. Keeping in view the economic importance of this fruit, the whole-genome sequencing was chosen to identify different novel genetic markers of economical important traits.

The advancement in next-generation sequencing technology, enabled the scholars to developed and screened different types of genomic molecular markers and applied for comparative studies, including SNP, InDel, and SSR. SSR are PCR based markers applied for population analysis, QTL mapping and genetic identification (Chen et al. 2008; Ollitrault et al. 2012). SNPs are considered as high frequencies markers belongs to last-generation molecular markers, well studied in both simple and complex organisms including plant and animal genomes. Researches are trying to develop more efficient ways to develop SNP markers as they enhance effectiveness of genotype analysis to tenfold. These are frequently utilized for the *in-silico* analysis of sequencing reads by mapping to a reference genome (Oueslati et al. 2017; Imai et al. 2018). For the present research we used GATK4 variant calling pipeline to identify the potential genetic markers from the sequence reads of *Citrus reticulata* by mapping on reference genome *Citrus clementina*. Similar studies were conducted in the regional countries like China, India, Japan, Honk Kong and Brazil. In China the comparative study of whole genome genetic variations between *Citrus clementina* and *Citrus trifoliata* ascertained a total of 3,062,798 and 7,744,203 SNPs. Moreover, a total of 252,679 InDels were identified in *Citrus clementina* and 747,897 in *Citrus trifoliata* against reference genome *Citrus clementina* (Liu et al. 2018). While in the current research we identified almost the same ratio of SNPs 3,503,033 and 656,370 Indels in *Citrus reticulata* ‘Kinnow’ indicating that many quantitative trait loci in plants have been linked to SNPs and a single SNP can have a significant phenotypic impact on plant (Nuñez et al. 2019). One of another previous work on Citrus reported the greatest number of SNPs in three citrus cultivars *Poncirus trifoliata, Citrus unshiu* and *Ponkan* with an average of one SNP after every 64 bp, 164 bp and one SNP every 1,002 bp respectively (Novelli et al. 2004) while the genome of *Citrus reticulata* is identified as 1 variant after every 69 bases.

It is observed that the true to type citrus trees showed the less number of SNPs in coding regions as compared to other plants like *Eucalyptus* species which showed 1.5 times higher SNPs per kb in coding regions. (Külheim et al. 2009). Similarly, the genomic coding regions of *Populus tremula, and* maize (Yamasaki et al. 2005) showed 16·7 and 23·25 SNPs per kb respectively. The previous studies on three principle species of Citrus reported various SNPs frequencies 15.15 SNPs per kb in *Citrus reticulata*, 4.70 in *Citrus maxima* and 2.21 in *Citrus medica* while the present study of *Citrus reticulata* showed 11.67 SNPs per kb which is 3.48 times less than the previous one (Garcia *et al*..2013). The SNP frequency acknowledge the ancestry of Kinnow and Clementine as Kinnow is the cross of King × Willowleaf while Clementine is a cross of Willowleaf × Sweet orange. King likely has sweet orange in its ancestry. It could be somewhat inbred. Anyway, the studying genome is not a wild type. All of these mandarins have considerable introgression of segments from pummelo which creates regions of high heterozygosity compared to regions of mandarin only origin (Ollitrault et al. 2011; Gill et al. 2022).

The genomic variants ultimately resulted in transition or transversion changes. Previously in satsuma observed more transition events as compared to transversion as 47.9% 36.1% respectively similarly the present research also showed higher transition as compared to transversion i.e., 65.44% and 34.55% respectively. Overall, G/A mutations were observed to be more as compared to C/T as compared to Satsuma where C/T mutations were common (Jiang et al. 2010). Another similar study identified the higher non-synonymous than synonymous changes in *Citrus reticulata* at the F3′H locus Ts:Tv=1:7 and HYB locus Ts:Tv=1:4 predicted the *Citrus reticulata* as more diversified taxa (Garcia et al. 2012, 2013) whereas we found 2,242,354 transitions (Ts) and 1,183,807 transversions (Tv) mutations, with a Ts:Tv ratio of 1:8 in *Citrus reticulata* genome. Estimation of the Ts/Tv provides insight into the process of molecular evolution and further help to model the evolutionary algorithms to have insight of the sequences of diverging genomes. Our studied citrus genome representing more transitions as compare to transversion which depicts its steady state evolutionary rate.

SNPs rates is directly proportional to the level of diversity, higher the number of SNPs greater the diversity among the citrus species, same as observed in other plants like rice (Terol et al. 2008; Mattia et al. 2022), Accordingly we hypothesized that the genetic variations reported in *Citrus reticulata* (present study) might be lower than that of *Citrus sinensis*, since *Citrus reticulata* is native to south Asia and most arose as crosses. Within a group such as satsumas there are many varieties that arose as mutants, but *Citrus reticulata* (broadly defined) includes many sexually produced hybrids. *Citrus reticulata* is the ancestral type, not *Citrus sinensis* so *Citrus reticulata* has a much longer evolutionary history, although both are hybrids (Wu et al. 2021). This diversity allowed the citrus growers in successful development of groups with different genotypes including blood oranges, navel oranges, and Valencia oranges. These aforementioned groups and few varieties within them were developed by breeders. They originated as mutants selected by growers, not breeders.

The finest alternative to SNP and SSR markers is InDel markers due to its simplicity, polymorphic nature, high transferability, efficient experimental procedure, low cost of development and richness across genome (Mills et al. 2006; Gao et al. 2012; Oliveira et al. 2022; Huang et al. 2022). These markers are also well studied to understand the phylogenetic relationship between closely related organisms (animals, plants, insects, bacteria) due to its ability of sequence divergence specifically from coding regions (Britten et al. 2003). In plants for marker-assisted breeding programs (Raman et al. 2006; Tiwari et al. 2022; Noda et al. 2022) and QTL mapping (Vasemäg et al. 2010; Goswami et al. 2022) many potential InDels markers had been applied due to genomic functional diversity. Moreover, to study the maternal inheritance in citrus many mitochondrial (Froelicher et al. 2011; Wang et al. 2022) and chloroplast (Caballero et al. 2015) InDels markers have been developed and studied. Citrus crops are diploid and highly heterozygous with smaller genomes (∼<400 Mb) have great primacy for InDel markers as compared to the other crops like maize due to its complex and large genome size. In this research, we developed 323,287 Ins and 333,083 Del, most of the InDels were reported form non-coding regions, almost same type of results was observed from different studies on Citrus (Fang et al. 2018), maize (Ching et al. 2002), and brassica (Park et al. 2010). InDel markers have not proven as broadly useful as SNPs and, other than small indels, are no longer widely used. Those in repeated sequences such as SSRs are not easily studied in large numbers and are limited by their rapid rate of evolution which leads to homoplasy (Ramadugu et al. 2013; Ollitrault et al. 2015; Ghada et al. 2019; Barbhuiya et al. 2022).

Codon changes can alter the amino acid content of proteins similarly in our protein such as GAC and GAT codes for Aspartate which is in green plants involved in the biosynthesis of different metabolites that play key roles in plants reproduction, development, growth and defense pathways (Torre et al. 2014). TCG and TCA both codes for the same amino acid i.e., Serine which is known as to play a fundamental role in cell signaling pathways (Ros et al. 2014). AAC and AAT codes for Asparagine, the very first amino acid identified about 200 years ago from plants (Vauquelin and Robiquet. 1806), involved in nitrogen storage in different tissues, and also well studied for its stress response under environmental conditions (Lea et al. 2007). AAG and AAA codes for Lysine an important role in plant growth and early plant yield (Galili 2002). Besides Lysine, Isoleucine, Glutamic acid, Histidine, and Tryptophan also well known to play a key role in crop growth. Tyrosine, Threonine, Cysteine known to have a tolerance against different plant diseases. Proline and Arginine well studied to have tolerance against hard environmental conditions like drought, heat, salinity, and frost. Glycine and alanine are important for chlorophyll formation in plant and ultimately photosynthesis. Methionine is responsible for fruit ripening while valine is responsible for seed production (Baqir et al. 2019).

Furthermore, study from China was conducted on Tangor (hybrid of two cultivars *Citrus sinensis* and *Citrus reticulata*) to identify the genes involved in the metabolism of starch, sucrose, cell wall, essential hormone, and phenyl propanoid pathways. These metabolic activities played a key role in fruit development and ripening, these genes are FLA4, FLA12, FT1, NADP-ME3, ALDH7B4, CA2, XTH9, EXPA5, LAC11, AAO, 5MAT, AMY1, BAM1, CT-BMY, BMY5, BAM1, PHO1, PHS2, SEX1, PWD, DPE2, PTPKIS1, BMY3, AMY3, GWD3, CCD7, NCED4, ABA1, AAO3, CCD1, NCED4, AAO1, HVA22J, SIR3 (Bi et al. 2022). All aforementioned genes are also identified in the present study with SNPs, InDels, MNPs affecting different genomic locations including CDS, Exon, Intron, UTRs produced high, low, moderate and modifier impact in genome. However, mechanism of action of the all the above-mentioned identified genes have yet not been verified and can be the topic of future research. Moreover, the interactions and function between these genes can be further studied during fruit development.

Despite the fact that most of the identified SNPs from present genome are not found within genes, but their prevalence and sturdiness highlight them as a valuable source of genomic DNA diversity for citrus breeding programs who are looking to develop improved and superior cultivars. Further, InDels are also known to cause phenotypic variations among the different species of Citrus. The presence of Indels can change the translation reading frame of genome.

## Conclusion

All the identified genetic markers in the present research can provide basic data to the researcher, academicians, farmers, breeders and policy makers for identification and improvement of subject plant by employing this preliminary genomic architecture insight of this valued plant, which might help them to improve growth, yield, shelf-life and nutritive value of the Kinnow fruit. Furthermore, targeted breeding programs may also be tailored to propagate its desired vested traits to improve the overall quality of the other members of the *Rutaceae* family to lift the country economy.

## Credit Author Statement

Sadia Jabeen (SJ) performed sample collection, wet-lab experimentation, WG variant calling pipeline, methodology/ data analysis and initial draft write-up. Rashid Saif (RS), WG variant calling pipeline design, data analysis, interpretation of results, initial draft write-up. Shagufta Naz (SN) conceptualization of the project, reviewing, proof reading and overall supervision. Rukhama Haq (RH) conceptualization of the project, co-supervision of wet-lab experimentation, reviewing and editing the draft. Akber Hayat (AH) identification of plants, reviewing and proof reading of the manuscript.

## Availability of Data

NGS-binary alignment maps (bam) files are submitted at NCBI SRA project ID: PRJNA821664. Furthermore, three supplementary tables and three figures are also provided along with the manuscript.

## Acknowledgement

Authors are obliged to the Emeritus Prof. Dr. Mikeal Roose, University of California, Riverside, USA and Dr. Subhas Hajeri, Director-Citrus Pest Detection Program, Tulare, CA, USA for reviewing and providing the valued suggestions and recommendation to improve the manuscript.

## References

Ahmed AH, Abd Ehhm (2003) Growth, uptake of some nutrients and productivity of Red Roomy vines as affected by spraying of some amino acids, magnesium and boron. Minia J of Agric Res Develop 23:649–666.

Ahmed K (2020) Third of Pakistan’s 2020 mandarin exports earmarked for the Middle East. Arab news PK https://arab.news/y4hrw. Accessed 6 December 2022

Altaf S, Khan MM, Jaskani MJ, Khan IA, Usman M, Sadia B, Awan FS, Ali A, Khan AI (2014) Morphogenetic characterization of seeded and seedless varieties of Kinnow Mandarin (‘Citrus reticulata’Blanco). Aust J Crop Sci 8:1542–1549. https://www.researchgate.net/publication/268152281

Baqir HA, Zeboon NH, Al-behadili Ajp (2019) The role and importance of amino acids within plants: a review. Plant Arch 19(2):1402–1410.

Barbhuiya AR, Khan ML, Dayanandan S (2022) Molecular phylogeny of Citrus species in the Eastern Himalayan region of Northeast India based on Chloroplast and Nuclear DNA sequence data. In: Kumar A, Choudhury B, Dayanandan S, Khan ML (eds) Molecular genetics and genomics tools in biodiversity conservation. Springer, Singapore,pp 185–201. https://doi.org/10.1007/978-981-16-6005-4-9

Bi X, Liao L, Deng L, Jin Z, Huang Z, Sun G, Xiong B, Wang Z (2022) Combined Transcriptome and Metabolome Analyses Reveal Candidate Genes Involved in Tangor (Citrus reticulata × Citrus sinensis) Fruit Development and Quality Formation. Int J Mol Sci 23:5457. https://doi.org/10.3390/ijms23105457

Bohry D, Ramos HCC, Santos PHD, Boechat MSB, Arêdes Fas, Pirovani AAV, Pereira MG (2021) Discovery of SNPs and InDels in papaya genotypes and its potential for marker assisted selection of fruit quality traits. Sci Rep 11:1–8. https://doi.org/10.1038/s41598-020-79401-z

Britten RJ, Rowen L, Williams J, Cameron RA (2003) Majority of divergence between closely related DNA samples is due to indels Proc Natl Acad 100:4661–4665. https://doi.org/10.1073/pnas.0330964100

Caballero JC, Alonso R, Ibañez V, Terol J, Talon M, Dopazo J (2015) Phylogenetic analysis of 34 Chloroplast genomes elucidates the relationships between wild and domestic species within the genus Citrus. Mol Bio and Evo 8:2015–2035. https://doi.org/10.1093/molbev/msv082

Chen C, Bowman KD, Choi YA et al (2008) EST-SSR genetic maps for Citrus sinensis and Poncirus trifoliata. Tree Genet Genomes 4:1–10. https://doi.org/10.1007/s11295-007-0083-3

Ching A, Caldwell KS, Jung M et al (2002) SNP frequency, haplotype structure and linkage disequilibrium in elite maize inbred lines BMC Genet 3:1–14. https://doi.org/10.1186/1471-2156-3-19

Cingolani P, Platts A, Wang LL, Coon M, Nguyen T, Wang L, Land SJ, Lu X, Ruden DM (2012) A program for annotating and predicting the effects of single nucleotide polymorphisms, SnpEff. Fly 6:80–92. https://doi.org/10.4161/fly.19695

Distefano G, Las CG, Deng X, Chai L (2020) Citrus reproductive biology from flowering to fruiting. In: Gentile A, La MS, Deng Z (eds) The citrus genome. Compendium of Plant Genomes. Springer, Cham, pp 167–176. https://doi.org/10.1007/978-3-030-15308-3-9

Duru S, Hayran S, Gül A (2022) The analysis of competitiveness of Mediterranean countries in the world citrus trade. Mediterr Agric Sci 35:21-26.10. https://doi.org/29136/mediterranean.1012466

Etxeberria E, Gonzalez P, Achor D, Albrigo G (2009) Anatomical distribution of abnormally high levels of starch in HLB-affected Valencia orange trees. Mol Plant Pathol 74:76–83. https://doi.org/10.1016/j.pmpp.2009.09.004

Fang Q, Wang L, Yu H et al (2018) Development of species-specific InDel markers in Citrus. Plant Mol Biol Rep 36: 653–662. https://doi.org/10.1007/s11105-018-1111-1

Fernandez CT (2022) Making a Pangenome Using the Iterative Mapping Approach. In: Edwards D (ed) Plant Bioinformatics. Methods in Molecular Biology. Humana, New York, pp 259–271. https://doi.org/10.1007/978-1-0716-2067-0-14

Froelicher Y, Mouhaya W, Bassene, JB et al (2011) New universal mitochondrial PCR markers reveal new information on maternal citrus phylogeny Tree Genet Genomes 7: 49–61. https://doi.org/10.1007/s11295-010-0314-x

Galili G (2002) New insights into the regulation and functional significance of lysine metabolism in plants. Annu Rev Plant Biol 53: 27.

Gao Q, Yue G, Li W, Wang J, Xu J, Yin Y (2012) Recent progress using High-throughput sequencing technologies in plant molecular breeding. J Integr Plant Biol 54:215–227. http://www.jipb.net, http://www.wileyonlinelibrary.com/journal/jipb

Garcia LA, Curk F, Snoussi TH et al (2013) A nuclear phylogenetic analysis: SNPs, indels and SSRs deliver new insights into the relationships in the ‘true citrus fruit trees’ group (Citrinae, Rutaceae) and the origin of cultivated species. Ann Bot 111:1–19. https://doi.org/10.1093/aob/mcs227

García-Lor A, Luro F, Navarro L et al (2012) Comparative use of InDel and SSR markers in deciphering the interspecific structure of cultivated citrus genetic diversity: a perspective for genetic association studies. Mol Genet Genomics 287:77–94. https://doi.org/10.1007/s00438-011-0658-4

Ghada B, Amel O, Aymen M et al (2019). Phylogenetic patterns and molecular evolution among ‘True citrus fruit trees’ group (Rutaceae family and Aurantioideae subfamily). Sci Hortic 253:87–98. https://doi.org/10.1016/j.scienta.2019.04.011.

Gill K, Kumar P, Kumar A et al (2022) Comprehensive mechanistic insights into the citrus genetics, breeding challenges, biotechnological implications, and omics-based interventions. Tree Genet Genomes 18:1–26. https://doi.org/10.1007/s11295-022-01544-z

Goswami M, Attri K, Goswami I (2022) Applications of molecular markers in fruit crops: A review. Int J Econ Plants 9:121–126.

Goto S, Yoshioka T, Ohta S, Kita M, Hamada H, Shimizu T (2016) Segregation and heritability of male sterility in populations derived from progeny of Satsuma mandarin. PLoS One 11:e0162408. https://doi.org/10.1371/journal.pone.0162408

Hayat F, Nawaz KM, Zafar SA, Balal R et al (2017) Surface Coating and Modified Atmosphere Packaging Enhances Storage Life and Quality of ‘Kaghzi lime’. J Agric Sci Technol 19:1151–1160. http://jast.modares.ac.ir/article-23-863-en.html

Huang X, Wu W, Su L, Lv H, et al (2022) Development and application of InDel markers linked to fruit-shape and peel-colour genes in Wax Gourd. Genes 13: 1567. https://doi.org/10.3390/genes13091567

Hughes HK, Rowland ME, Onore CE et al (2022) Dysregulated gene expression associated with inflammatory and translation pathways in activated monocytes from children with autism spectrum disorder. Transl Psychiatry, 12:1–9. https://doi.org/10.1038/s41398-021-01766-0

Imai A, Nonaka K, Kuniga T et al (2018) Genome-wide association mapping of fruit-quality traits using genotyping-by-sequencing approach in citrus landraces, modern cultivars, and breeding lines in Japan. Tree Genet Genomes 14: 24. https://doi.org/10.1007/s11295-018-1238-0

Jaskani MJ, Kwon SW, Kim DH (2005) Comparative study on vegetative, reproductive and qualitative traits of seven diploid and tetraploid watermelon lines. Euphytica 145:259–268. https://doi.org/10.1007/s10681-005-1644-x

Jiang D, Ye QL, Wang FS, Li CAO (2010) The mining of citrus EST-SNP and its application in cultivar discrimination. Agricl Sci China, 9:179–190.https://doi.org/10.1016/S1671-2927(09)60082-1

Jiang P, Zhu T, Liu J, Tao X, Xue Z et al (2022) Mitochondrial DNA variant spectrum and the association with chronic tic disorders. Eur J Neurol 29:3187–3196.

Kamal GM, Anwar F, Hussain AI, Sarri N, Ashraf MY (2011) Yield and chemical composition of Citrus essential oils as affected by drying pretreatment of peels. Int Food Res J 18:1275–1282.

Khan IA, Kender WJ (2007) Citrus breeding: Introduction and objectives. Citrus genetics, breeding and biotechnology. CABI. United Kingdom.

Külheim C, Hui YS, Maintz J, Foley WJ, Moran GF (2009) Comparative SNP diversity among four Eucalyptus species for genes from secondary metabolite biosynthetic pathways. BMC genom 10:1–11. https://doi.org/10.1186/1471-2164-10-452

Lea PJ, Sodek L, Parry MA, Shewry PR, Halford NG (2007) Asparagine in plants. Ann of Appl Biol150:1–26. https://doi.org/10.1111/j.1744-7348.2006.00104.x

Li H, Durbin R (2010) Fast and accurate long-read alignment with Burrows–Wheeler transform. Bioinformatics, 26:589–595.https://doi.org/10.1093/bioinformatics/btp698

Liu E, Li J, Ou S, Dong B, Yang B, Zhou Y (2022) The complete mitochondrial genome of Semblis atrata (Trichoptera: Phryganeidae). Mitocho DNA Part B, 7:6, 956–958. https://doi.org/10.1080/23802359.2022.2080595

Liu TJ, Zhou JJ, Chen FY, Gan Z-M, Li Y-P, Zhang J-Z, Hu C-G (2018) Identification of the Genetic Variation and Gene Exchange between Citrus Trifoliata and Citrus Clementina. Biomolecules 8:182. https://doi.org/10.3390/biom8040182

Madian AM, Refaai MM (2011) The synergistic effects of using B vitamins with the two amino acids tryptophan and methionene in Thompson seedless grapevines. Minia J Agric Res Develop 3:445–455.

Marbouty M, Koszul R (2022) Metagenomes Binning Using Proximity-Ligation Data. In: Bicciato S, Ferrari F (eds) Hi-C Data Analysis. Methods in Molecular Biology. Humana, New York, pp 163–181 https://doi.org/10.1007/978-1-0716-1390-0-8

Mattia MR, D. D, Yu Q, Kahn T, Roose M, Hiraoka Y, Wang Y, Munoz P, Gmitter FG Jr (2022) Genome-Wide Association Study of Healthful Flavonoids among Diverse Mandarin Accessions. Plants 11:317. https://doi.org/10.3390/plants11030317

Mills RE, Luttig CT, Larkins CE, Beauchamp A, Tsui C, Pittard WS, Devine SE (2006) An initial map of insertion and deletion (INDEL) variation in the human genome. Genome Res 16(9):1182-1190.1182-1190.https://doi.org/10.1101/gr.4565806

Montalt R, Vives MC, Navarro L, Ollitrault P, Aleza P (2021) Parthenocarpy and self-incompatibility in Mandarins. Agronomy 11:2023.https://doi.org/10.3390/agronomy11102023

Murray MG, Thompson WF (1980) Rapid isolation of high molecular weight plant DNA. Nucleic Acids Res 8:4321–4326. https://doi.org/10.1093/nar/8.19.4321

Naqvi SAH, Wang J, Malik MT, Umar UUD, Hasnain A, Sohail MA, Shakeel MT, Nauman M, Hassan MZ, Fatima MJA (2022) Citrus Canker-distribution, taxonomy, epidemiology, disease cycle, pathogen biology, detection, and management: A critical review and future research agenda. Agronomy 12(5):1075. https://doi.org/10.3390/agronomy12051075

Naz S, Shahzadi K, Rashid S, Saleem F, Zafarullah A, Ahmad S (2014) Molecular characterization and phylogenetic relationship of different citrus varieties of Pakistan. J Anim Plant Sci 24:315–320

Noda T, Daiou K, Mihara T et al (2022) Efficient method for generating citrus hybrids with polyembryonic Satsuma mandarin as the female parent. Mol Breed 42:1–15. https://doi.org/10.1007/s11032-022-01324-6

Novelli VM, Takita MA Machado MA (2004) Identification and analysis of single nucleotide polymorphisms (SNPs) in citrus. Euphytica 138:227–237. https://doi.org/10.1023/B:EUPH.0000047086.47988.82

Nuñez LG, Balladares C, Pavez C, Urra C, Sanhueza D, Vendramin E et al (2019) High-density genetic map and QTL analysis of soluble solid content, maturity date, and mealiness in peach using genotyping by sequencing, Sci Hortic 257:108734. https://doi.org/10.1016/j.scienta.2019.108734.

Oliveira M, Azevedo L (2022) Molecular Markers: An Overview of Data Published for Fungi over the Last Ten Years. J Fungi 8:803. https://doi.org/10.3390/jof8080803

Ollitrault F, Terol J, Martin AA, Pina JA, Navarro L, Talon M, Ollitrault P (2012) Development of indel markers from Citrus clementina (Rutaceae) BAC-end sequences and interspecific transferability in Citrus. Am J Bot 99:e268–e273. https://doi.org/10.3732/ajb.1100569

Ollitrault P, Garcia-Lor A, Terol J, Curk F, Ollitrault F, Talón M, Navarro L (2015) Comparative values of SSRs, SNPs and InDels for citrus genetic diversity analysis. Acta Hortic 1065:457–466. https://doi.org/10.17660/ActaHortic.2015.1065.56

Ollitrault P, Terol JF, Chen C, Federici CT, et al (2011). A reference linkage Map of C. clementina based on SNPs, SSRs and indels:477. https://agritrop.cirad.fr/559981

Oueslati A, Salhi-Hannachi A, Luro F, Vignes H, Mournet P, Ollitrault P (2017) Genotyping by sequencing reveals the interspecific C. maxima / C. reticulata admixture along the genomes of modern citrus varieties of mandarins, tangors, tangelos, orangelos and grapefruits. PLoS ONE 12: e0185618. https://doi.org/10.1371/journal.pone.0185618

Panaro NJ, Yuen PKI, Sakazume T, Fortina P, Kricka LJ, Wilding P (2000) Evaluation of DNA Fragment Sizing and Quantification by the Agilent 2100 Bioanalyzer, Clin Chem 46:1851–1853. https://doi.org/10.1093/clinchem/46.11.1851

Park S, Yu HJ, Mun JH et al (2010) Genome-wide discovery of DNA polymorphism in Brassica rapa. Mol Genet Genomics 283:135–145. https://doi.org/10.1007/s00438-009-0504-0

Prasad H, Thakur M, Gupta AK, Prasad D (2015) Effect of Foliar Application of 2, 4-D, Urea and Zinc Sulphate on Fruit Drop, Yield and Fruit Quality of Kinnow Mandarin. Int j bio-resour stress manag 6.

Ramadugu C, Pfeil BE, Keremane ML, Lee RF, Maureira-Butler IJ, Roose ML (2013) A Six Nuclear Gene Phylogeny of Citrus (Rutaceae) Taking into Account Hybridization and Lineage Sorting. PLoS ONE 8: e68410. https://doi.org/10.1371/journal.pone.0068410

Raman H, Raman R, Wood R et al (2006) Repetitive Indel Markers within the ALMT1 Gene Conditioning Aluminium Tolerance in Wheat (Triticum aestivum L.). Mol Breed 18:171–183. https://doi.org/10.1007/s11032-006-9025-2

Raudvere U, Kolberg L, Kuzmin I, Arak T et al (2019) g:Profiler: a web server for functional enrichment analysis and conversions of gene lists (2019 update). Nucleic Acids Rese 47:W191–W198. https://doi.org/10.1093/nar/gkz369

Rehman A, Deyuan Z, Hussain I, Iqbal MS, Yang Y, Jingdong L (2018) Prediction of Major Agricultural Fruits Production in Pakistan by Using an Econometric Analysis and Machine Learning Technique, Int J of Fruit Sci 18:445–461.10.1080/15538362.2018.1485536

Ros R, Muñoz BJ, Krueger S (2014) Serine in plants: biosynthesis, metabolism, and functions. Trends Plant Scie 19:564–569.https://doi.org/10.1016/j.tplants.2014.06.003.

Sabir I (2010) Pakistan, the largest kinnow grower. Daily times. https://defence.pk/pdf/threads/pakistan-currently-the-largest-kinnow-grower.84281/

Sambrook J, Fritsch E, Maniatis T (1989) Molecular cloning: a laboratory manual. Cold Spring Harbor, New York

Sambrook j, Green MR (2012) Molecular cloning: A laboratory manual. Cold Spring Harbor, New York

Terol J, Naranjo MA, Ollitrault P et al (2008) Development of genomic resources for Citrus clementina: Characterization of three deep-coverage BAC libraries and analysis of 46,000 BAC end sequences. BMC Genom 9:1–12. https://doi.org/10.1186/1471-2164-9-423

Tiwari JK, Yerasu SD, Rai N, et al (2022) Progress in Marker-Assisted Selection to Genomics-Assisted Breeding in Tomato. Crit Rev Plant Sci 41:321–350. https://doi.org/10.1080/07352689.2022.2130361

Torre FDL, Cañas RA, Pascual MB, Avila C, Cánovas FM (2014) Plastidic aspartate aminotransferases and the biosynthesis of essential amino acids in plants. J Exp Bot 65:5527–5534. https://doi.org/10.1093/jxb/eru240

Usman M, Fatima B, Gillani KA, Khan MS, Khan MM (2008) Exploitation of potential target tissues to develop polyploids in citrus. Pak J Bot 40:1755–1766.

Vasemäg A, Gross R, Palm D et al (2010) Discovery and application of insertion-deletion (INDEL) polymorphisms for QTL mapping of early life-history traits in Atlantic salmon. BMC Genom 11:1–11. https://doi.org/10.1186/1471-2164-11-156

Vauquelin LN, Robiquet PJ (1806) The discovery of a new plant principle in Asparagus sativus. Ann Chim 57:14.

Wang N, Li C, Kuang L, Wu X et al (2022). Pan-mitogenomics reveals the genetic basis of cytonuclear conflicts in citrus hybridization, domestication, and diversification. Proc Natl Acad Sci 119:e2206076119. https://doi.org/10.1073/pnas.2206076119

Wu G, Prochnik S, Jenkins J et al (2014) Sequencing of diverse mandarin, pummelo and orange genomes reveals complex history of admixture during citrus domestication. Nat Biotechnol 32:656–662. https://doi.org/10.1038/nbt.2906

Wu GA, Sugimoto C, Kinjo H et al (2021) Diversification of mandarin citrus by hybrid speciation and apomixis. Nat Commun 12: 4377. https://doi.org/10.1038/s41467-021-24653-0

Wu G, Terol J, Ibanez V et al (2018) Genomics of the origin and evolution of Citrus. Nature 554:311–316. https://doi.org/10.1038/nature25447

Yamasaki M, Tenaillon MI, Vroh BI, et al (2005) A large-scale screen for artificial selection in maize identifies candidate agronomic loci for domestication and crop improvement. Palt Cell. 17:2859–2872. https://doi.org/10.1105/tpc.105.037242

Zhao C, Wang F, Lian Y, Xiao H, Zheng J (2020) Biosynthesis of citrus flavonoids and their health effects. Crit Rev Food Sci Nutr 60:566–583. https://doi.org/10.1080/10408398.2018.1544885

