## Supplementary Figure 1 for "Whole-Genome Sequencing and Variant Discovery of *Citrus reticulata* ‘Kinnow’ from Pakistan"

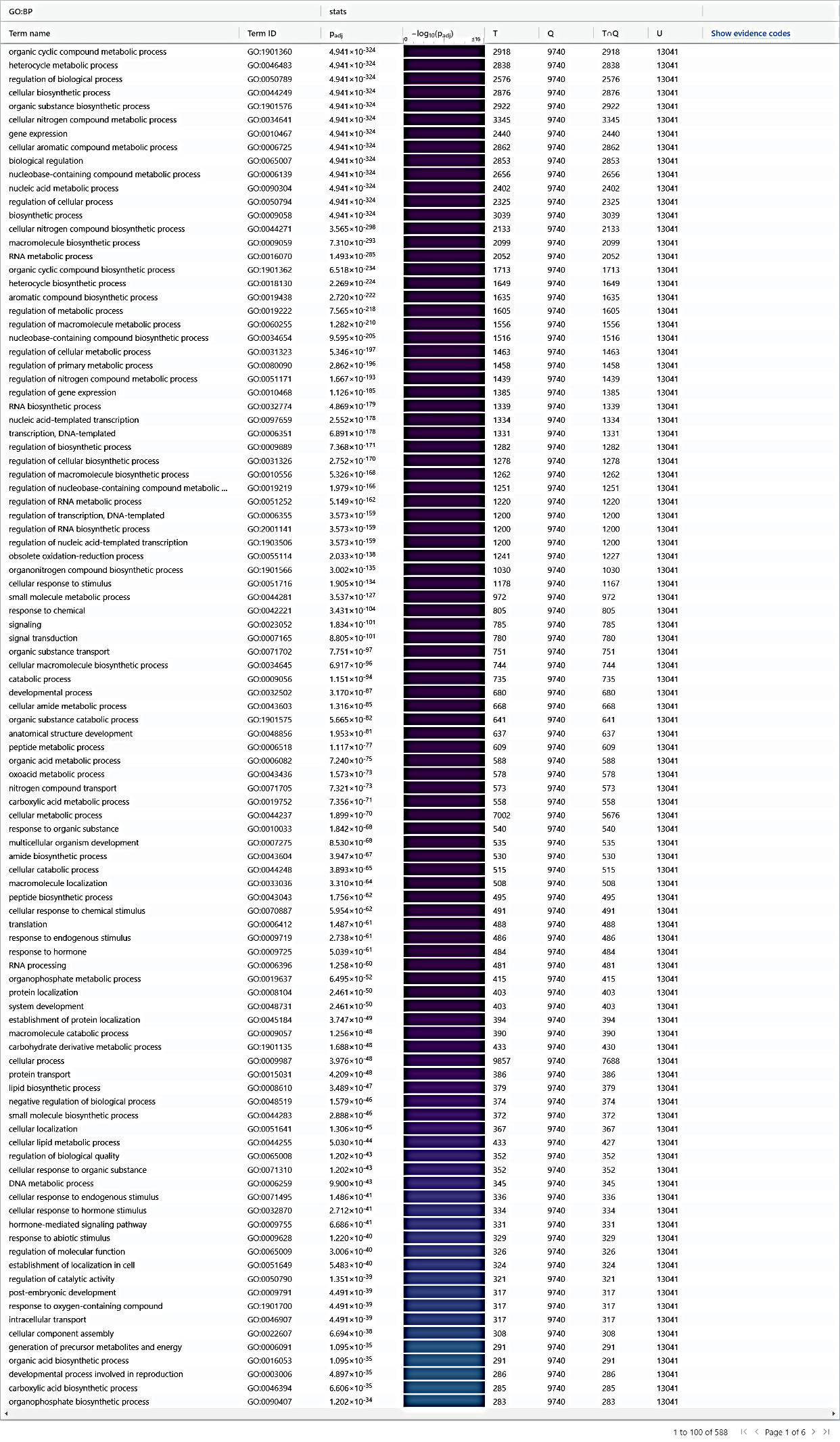

**Supplementary Figure S1.** Heatmap represents the annotated transcripts with term ids and adjusted *p*-value to different biological processes
