## Supplementary Figure 2 for "Whole-Genome Sequencing and Variant Discovery of *Citrus reticulata* ‘Kinnow’ from Pakistan"

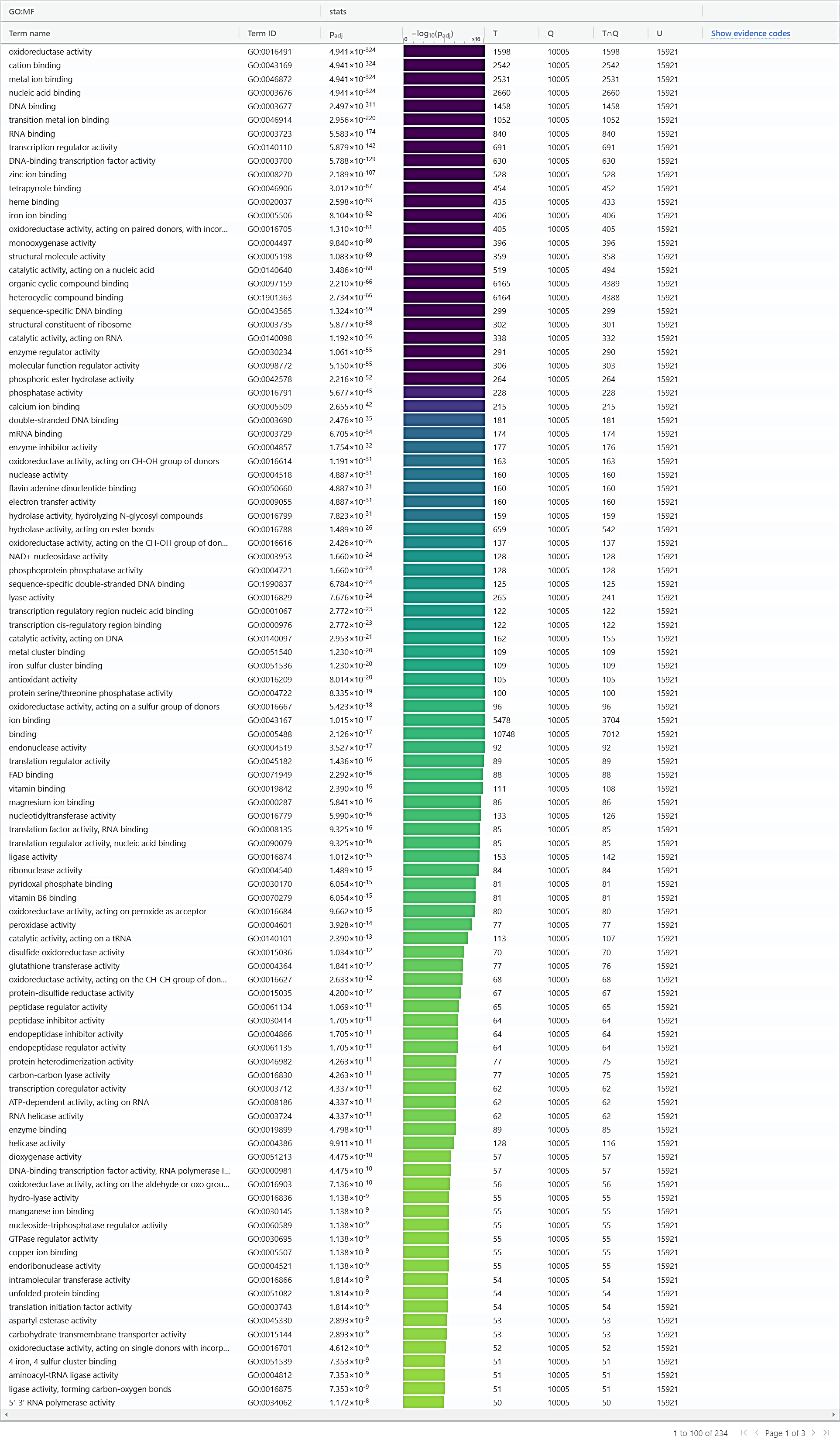

**Supplementary Figure. 2** heatmap represents the annotated transcripts with term ids and adjusted p value to different molecular functions
