## Supplementary Figure 3 for "Whole-Genome Sequencing and Variant Discovery of *Citrus reticulata* ‘Kinnow’ from Pakistan"

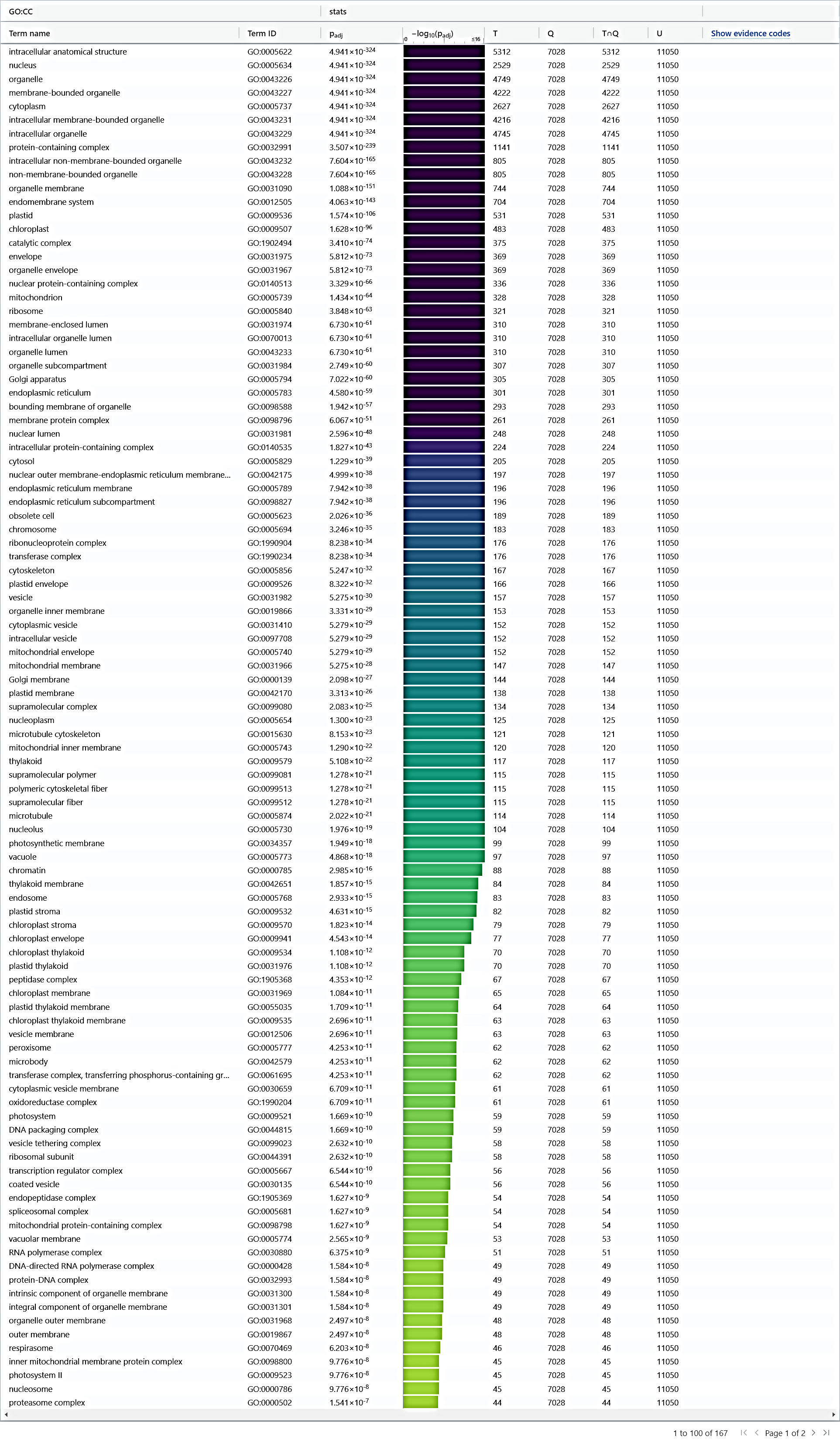

**Supplementary Figure. 3** heatmap represents the annotated transcripts with term ids and adjusted p value to different cellular components
